# Self-organized map: the new aproach for study of genetic divergence in kale

**DOI:** 10.1101/2020.05.14.095711

**Authors:** Orlando Gonçalves Brito, Valter Carvalho de Andrade Júnior, Alcinei Mistico Azevedo, Maria Thereza Netta Lopes Silva, Ludimila Geiciane de Sá, Clóvis Henrique Oliveira Rodrigues, Ana Clara Gonçalves Fernandes

## Abstract

The objective of this study was to study the genetic divergence between genotypes of kale, to propose a methodology for the use of neural networks of the SOM type and to test its efficiency through Anderson’s discriminant analysis. We evaluated 33 families of half-siblings of kale and three commercial cultivars. The design was a randomized block with four replications with six plants per plot. A total of 14 plant-level quantitative traits were evaluated. Genetic values were predicted at family level via REML / BLUP. For the study of divergence, neural networks of the SOM type (Self-organizing Map) were adopted. We evaluated different network architectures, whose consistencies of the clusters were identified by the Anderson discriminant analysis and by the number of empty clusters. After selecting the best network configuration, a dissimilarity matrix was obtained, from which a dendrogram was constructed using the UPGMA method. The best network architecture was formed with five rows and one column, totaling five neurons and consequently five clusters. The greatest dissimilarity was established between clusters I and V. The crossing between the genotypes of cluster I and those belonging to clusters III and V are the most recommended, since they aim to recombine families with characteristics of interest to the improvement and high dissimilarity. Anderson’s discriminant analysis showed that the genotype classification was 100% correct, indicating the efficiency of the methodology used.

## Introduction

Genetic improvement is one of the main tools to increase crop yield in a sustainable way [1]. However, for genotype selection in breeding programs, knowledge of genetic dissimilarity among the genotypes of the population to be improved is essential [2, 3]. Several molecular techniques have been used for genetic improvement. However, the use of phenotypic characters in the study of dissimilarity is more appropriate. This is due to its direct association with the attributes of interest in breeding. Thus, it is necessary to select dissimilar genotypes for recombination in order to provide greater genetic variability in the segregating populations [4].

Several studies on the evaluation of genetic divergence in brassicas have been carried out in different countries [5, 6, 7]. However, few studies have been done specifically with kale (*Brassica oleracea* var. *acephala*) [8].

Often genetic diversity studies have been performed using traditional multivariate techniques such as dendrograms, major components, and canonical variables. However, there is the possibility of carrying out these studies through computational intelligence using artificial neural networks (ANNs) [1, 9]. The main advantages of RNAs are their non-parametric approach, tolerance to data loss, and the need for detailed information about the modeling system as a design and genealogies [10, 11].

Self-organizing Map (SOM) neural networks are a type of exploratory multivariate analysis tool that allows, through artificial computational intelligence, to design high-dimensional data in a smaller dimensional space, without loss of information [12]. This new organization prioritizes maintaining the structure, such as clusters and information relationships [13]. This reinforces its constant use in several grouping and process characterization works. However, the network topology is usually selected subjectively. In addition, at the beginning of the iterative process of SOM networks, the synaptic weights are random, which can generate several results for the same data set and network configuration. This may lead to the discrediting of this methodology, which requires the implementation of strategies to mitigate this problem.

The objective of this study was to study the genetic divergence between families of half-siblings of leaf kale in order to select parents, to propose a new methodology for the use of SOM neural networks and to test its efficiency through Anderson’s discriminant analysis.

## Material and methods

### Experiment Implantation and data collection

The experiment was conducted at the Institute of Agricultural Sciences (ICA) of the Federal University of Minas Gerais, located in the municipality of Montes Claros-MG, from October 2016 to August 2017. The region is located at coordinates 16° 41′S and 43° 50′W and altitude of 646,29 m, with climate AW type (tropical savanna climate with dry winter and rainy summer) according to classification of Köeppen [14], and included in the region belonging to the Brazilian semiarid.

The experimental design was a randomized block design with four replications. The evaluated treatments consisted of 36 genotypes of kale, with 33 half-sib families and three commercial cultivars (Butter, Portuguese Butter and Georgia Butter). The plants of the half-sib families were obtained through cloning, using sprouts from an experiment conducted at the Federal University of the Jequitinhonha and Mucuri Valleys (UFVJM).

Sprouts were removed from the plants when they were approximately 5 cm. Subsequently, they were rooted in styrofoam trays of 72 cells filled with commercial substrate. The sprouts remained in a greenhouse for 40 days until they reached seedlings for planting. Due to the difficulty of obtaining sprouts in the commercial cultivars, the seedlings of the same ones were produced from seeds acquired in the local commerce. Both the seedlings of the half-sib families and the commercial cultivars were taken to the field when they presented the same pattern.

Soil preparation was carried out through plowing and two harrowing. Afterwards, beds of 1.20 m were prepared, with the aid of mechanized equipment. In the plot were planted six seedlings of the corresponding genotype. The spatial arrangement of the plants was in double row, spaced 1.00 × 0.50 m, representing a population of 20 thousand ha^−1^ plants. Planting fertilization and other cultural treatments such as irrigation, phytosanitary management and cover fertilization were carried out according to crop needs.

We evaluated 14 characteristics of quantitative origin in each plant of the plot (S1 Table). The number of shoots, leaves, the average mass of fresh leaf and the production of leaves were evaluated every 15 days. At 160 days the leaf length, limbus length, petiole length, petiole base diameter, petiole diameter and leaf width were evaluated. At 170 the plant height, stem height, stem base and middle were measured. These evaluation dates occurred in these periods (160 and 170 DAT) due to the fact that the plants are already considered adults and in full production.

### Data analysis

The genetic divergence between families was characterized at the family level. The statistical model 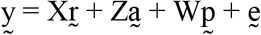, in which 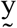 is the data vector, 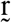 is the vector of the repetitive effects (assumed as fixed) added to the general average, 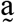 is the vector of the individual additive genetic effects (random), 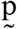 is the vector of the plots effects (random), 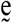 is the vector of errors or residuals (random). Capital letters represent the incidence matrices for these effects. For this, genetic-statistical software Selegen-REML / BLUP [15] was used. Subsequently, the data were standardized to obtain an average of zero and a standard deviation of one. Neural networks of the SOM type were adopted for the study. Different network architectures were evaluated, varying the number of rows from 1 to 6 and columns from 1 to 6. Excluding the combination of a row and a column, which would provide a single cluster, there were 35 tested configurations.

At the beginning of the iterative process of SOM networks, the synaptic weights are random, this can lead to several results for the same data set and network configuration. Therefore, the best network architecture was established from 1000 trainings performed for each of the 35 configurations. For each of them the average hit rate was estimated through Anderson’s discriminant analysis [16] and the smaller number of empty clusters. The best network architecture was selected from the highest average hit rate and the lowest number of empty clusters, simultaneously, following the parsimony principle.

Having determined the best configuration (topology) of the network, it was submitted to 1000 new trainings, and later a matrix of dissimilarity was built. The dissimilarity between each pair of genotypes was determined from the frequency at which they were allocated in different clusters.

Then the dendrogram of dissimilarity was made from the UPGMA method (Unweighted Pair Group Method using Arithmetic averages). The number of clusters of the dendrogram was established according to the number of neurons formed in the best network configuration. The consistency of the clustering was verified through Anderson’s discriminant analysis.

For the complementation of the clustering study, the graphical dispersion of the dissimilarity matrix in two-dimensional projection [17] was performed. In this procedure, the dissimilarity measures are converted into scores relative to two variables that, when represented in scatter plots, will reflect the distances originally obtained.

All analyzes were done in R [18] software (S2 Table). For the use of SOM networks, the RSNNS package was used. In order to obtain the dendrogram, the hclust function was used.

## Results

It was found that the best topology was established with five rows and one column (Fig 1A). This configuration was the one that allowed the Anderson Anderson discriminant analysis (98.72%) (Fig 1A) and showed a lower number of empty clusters (zero) (Fig 1B). Therefore, it was possible to form five neurons (five lines x one column), implying the formation of five clusters.

**Figure 1.**
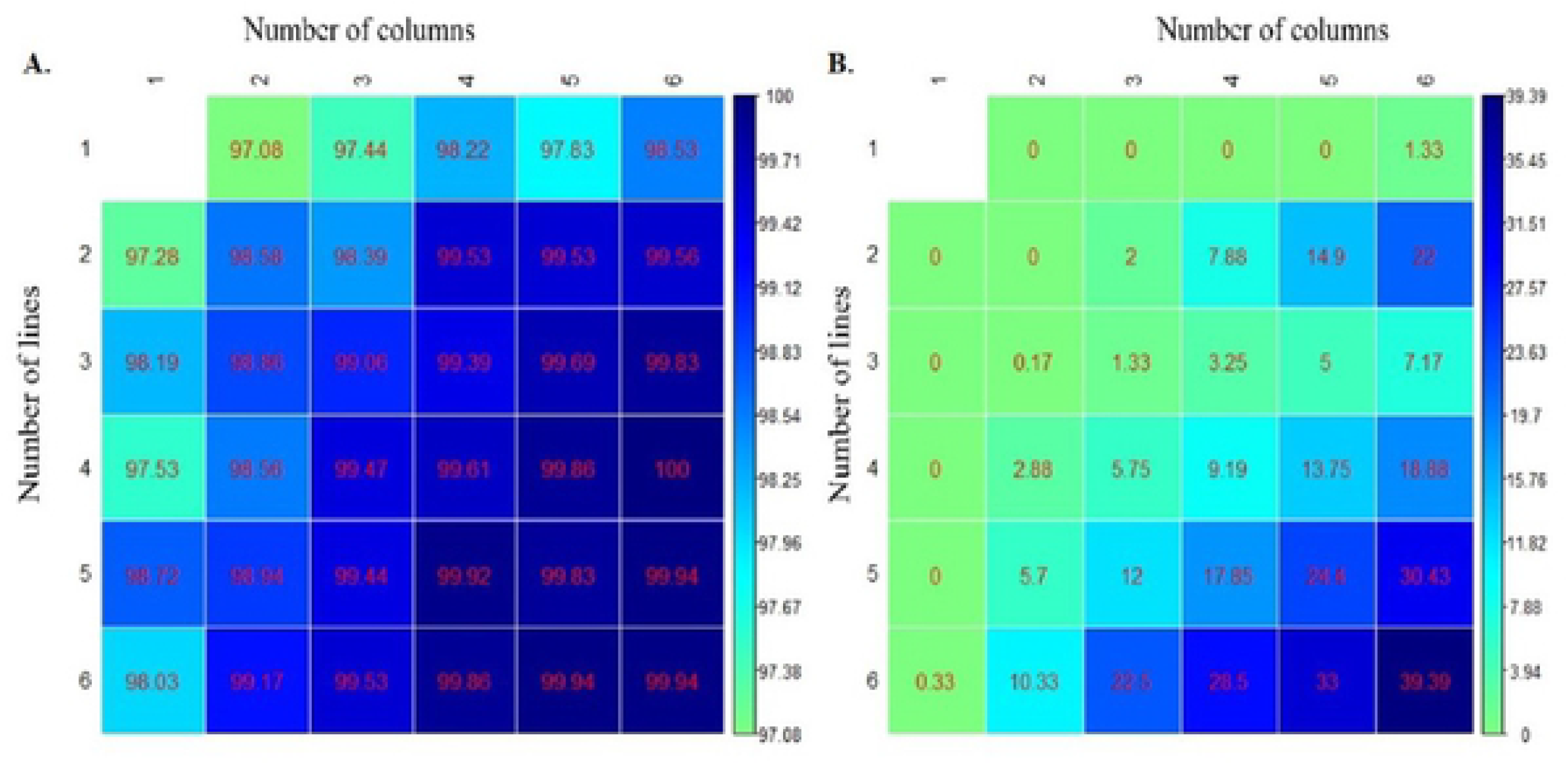
Average percentage of correct classifications by Anderson discriminant analysis (A) and percentage of empty clusters considering 35 SOM (B) network configurations.

From the best network architecture, the dissimilarity matrix was obtained from Kohonen’s self-organizing maps (Fig 2). Lighter colors indicate smaller genetic distances between genotypes, while darker colors represent greater distances, that is, greater genetic divergence. It was observed that there was high genetic divergence between most of the genotypes (Fig 2). Commercial cultivars (COM1, COM2 and COM3) were similar to each other but divergent in relation to most half-sib families. The families most genetically similar to the commercial cultivars were F8, F22 and F23.

**Figure 2.**
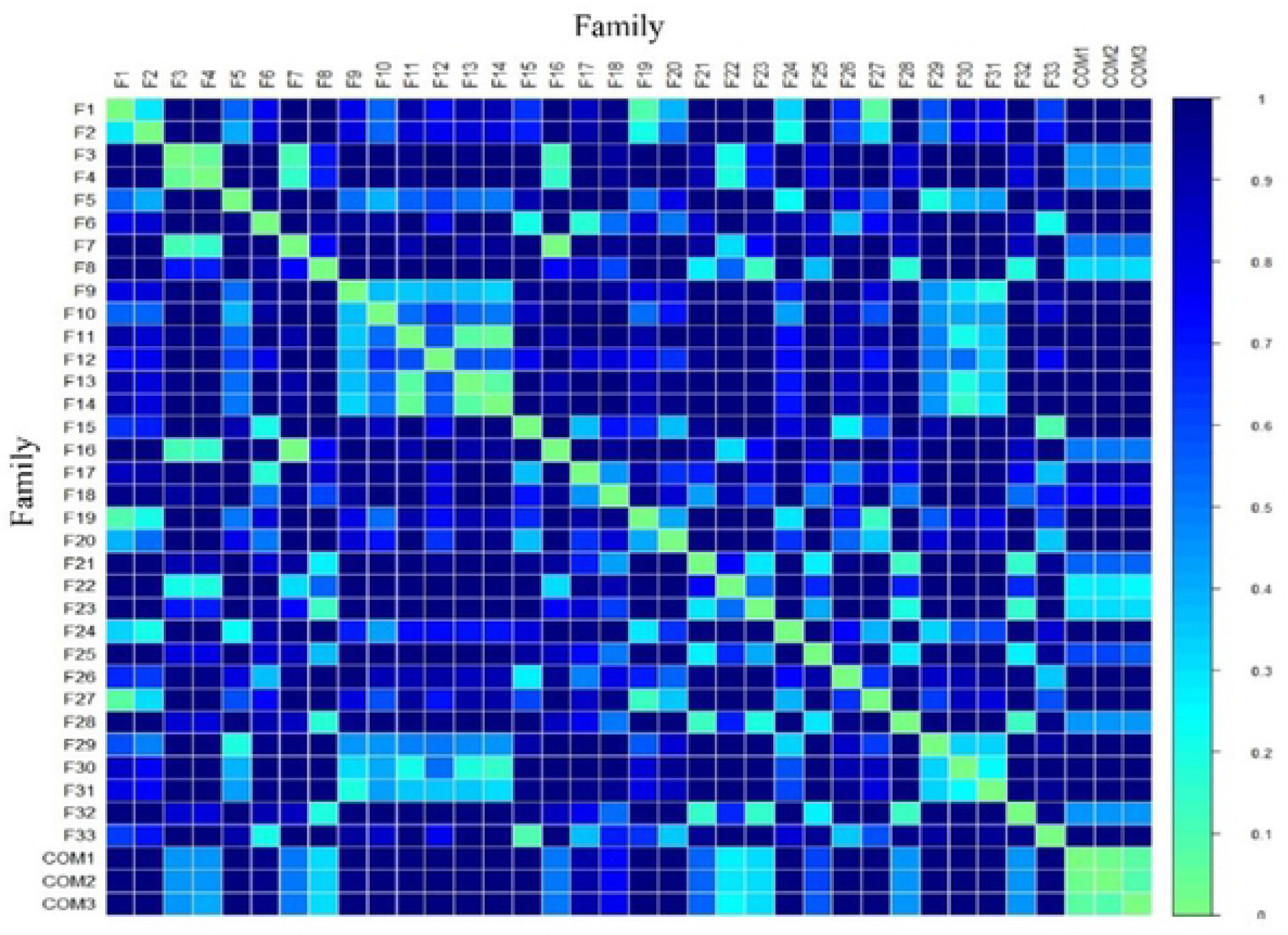
Graphical representation of the dissimilarity matrix obtained by Kohonen self-organizing maps. “F” refers to the half-sib family and “Com” to the commercial cultivar.

Analyzing the clusters formed by the UPGMA method (Fig 3), it is verified that the first cluster (I) was composed of five half-sib families (F3, F4, F7, F16 and F22) and the three commercial cultivars. These materials were characterized by low number of sprouts, leaves and yield, as well as intermediate values of the base and middle stem diameters (Fig 4).

**Figure 3.**
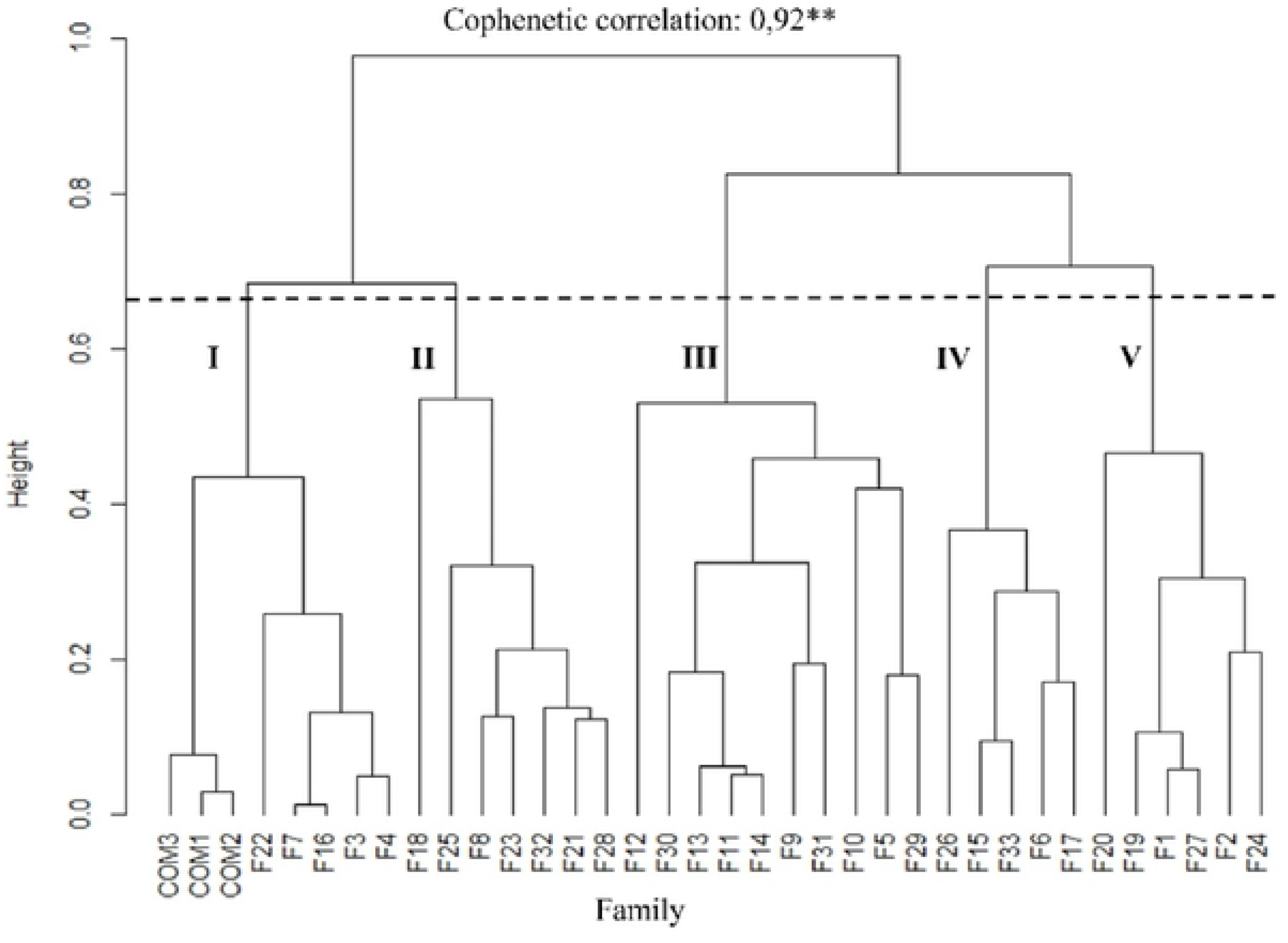
Dendrogram obtained by the UPGMA method from the dissimilarity matrix obtained by SOM type networks for the composition of the clusters (I, II, III, IV and V) considering 14 quantitative characteristics in leaf cabbage genotypes.

**Figure 4.**
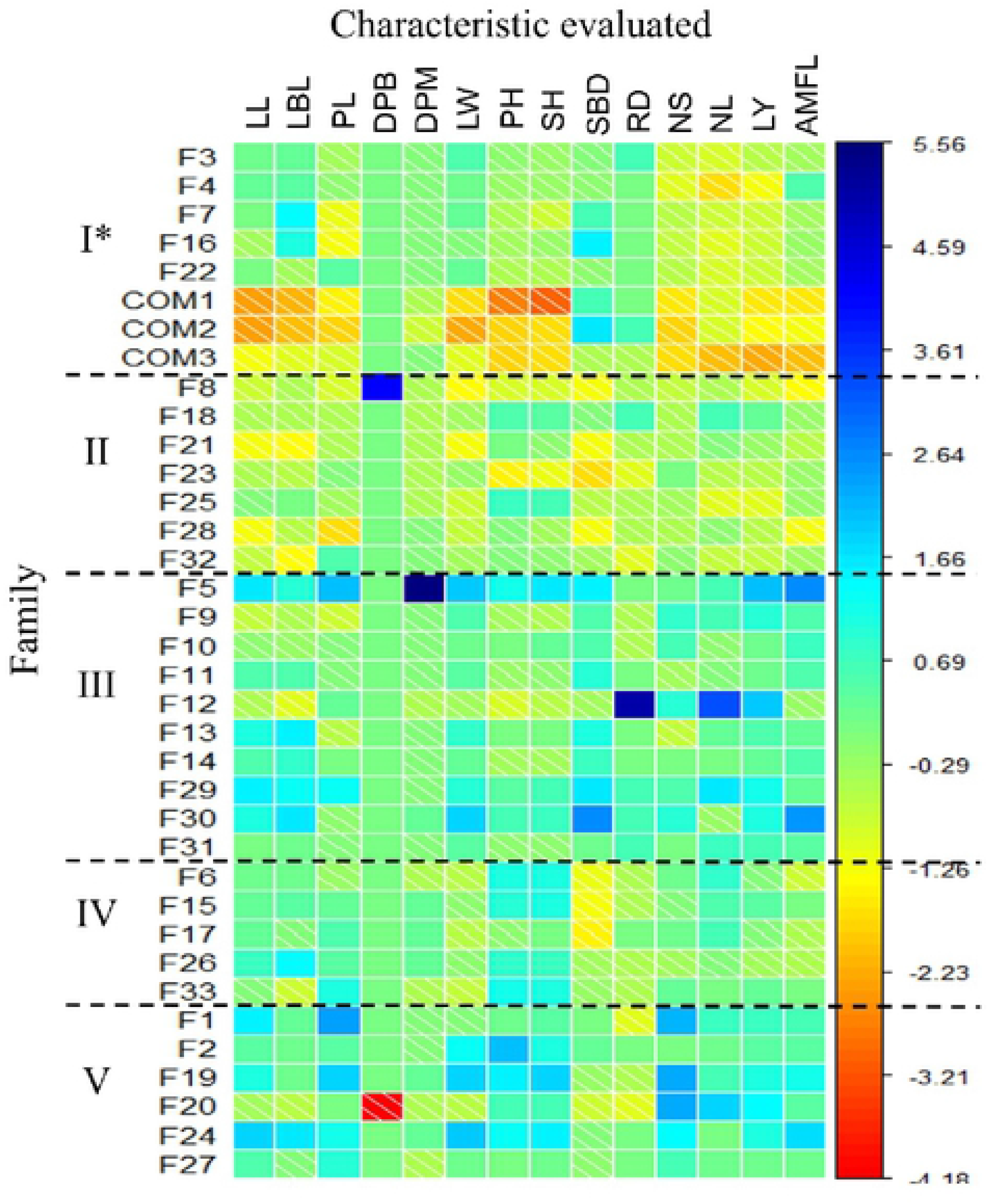
Graphical representation of the standardized genetic values for the study of the contribution of each characteristic to the composition of the clusters. Characteristics: NS: number of shoots; NL: number of leaves; AMFL: average mass of fresh leaf; LY: leaf yield; LL: leaf length; LBL: limbus length; PL: petiole length; DPB: diameter of the base of the petiole; DPM: diameter of the petiole medium; LW: leaf width; PH: plant height; SH: stem height; SBD: stem base diameter; RD: diameter of the base of the stem. * Cluster.

The commercial cultivars showed the lowest values for all the characteristics when compared with the half-sib families evaluated. This was only observed for the petiole diameters (DBP and DPM) and stem (SBD and RD). It was also observed that the diameter of the base of the petiole presented similar values for all families, except for the family F8 (higher value) and F20 (smaller value).

The second cluster (II) consisted of seven half-sib families (F8, F18, F21, F23, F25, F28 and F32). These families, in general, presented intermediate values for the number of sprouts and diameter of the middle of the petiole, in addition to plants with smaller stem base diameter and smaller width of leaves. As observed for cluster I, these families also presented lower agronomic performance than the other genotypes, with low leaf production (Fig 4). The third cluster (III) was formed by ten families of half-brothers (F5, F9, F10, F11, F12, F13, F14, F29, F30 and F31). This cluster was characterized by presenting families with higher genetic values for practically all the characteristics evaluated, especially for yield, number of shoots, stem base diameter and leaf width.

The fourth and fifth cluster (IV and V) were composed of five and six half-sib families, respectively. These clusters were composed of families with higher plant height and stem height, and cluster V also presented high number of shoots and high leaf yield. It should be emphasized that the families of clusters IV and V, also, have characteristics that are not of interest for the improvement of the crop, plants of low diameter of the bases of the stem (cluster IV), with high values for height of plants and petiole length (mainly in cluster V), as well as a high number of buds and leaves too large (high values of LL and LW) (Fig 4).

In a possible selection, clusters I, III and V should be prioritized, as they present half-sibling families with agronomic characteristics of interest for improvement, such as lower plant heights and lower number of shoots (cluster I), larger stem diameter (cluster III) and high leaf yield (cluster III and V). The recombination of these characteristics may contribute to a greater heterotrophic effect in the segregating population, which is of interest.

It was verified that the cluster IV was the most homogeneous, presenting smaller average genetic distance within the cluster (0.232) (Table 1). This can be verified through the smaller variation of color existing between the families within each assessed characteristic (Fig 4). The fact that this cluster presents few genotypes may also have contributed to this greater homogeneity.

**Table 1.**
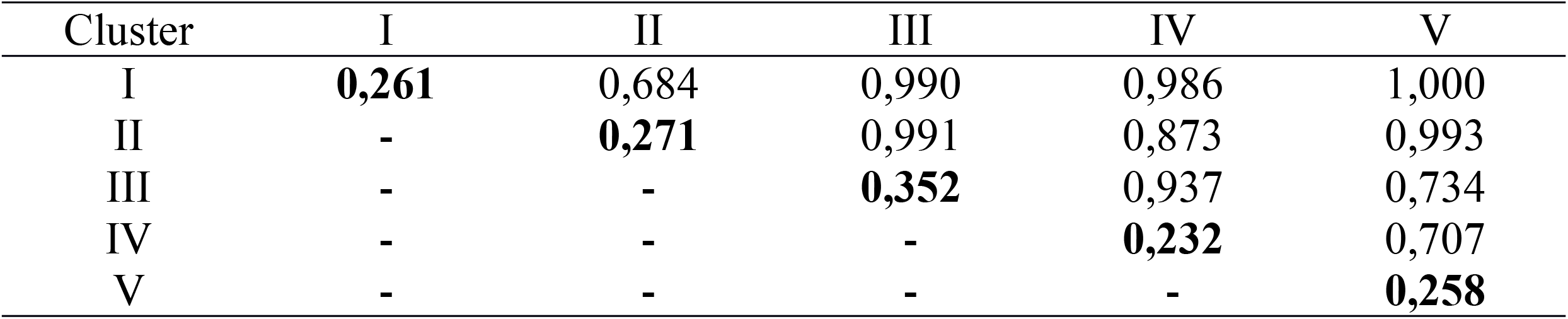
Average distances within (main diagonal in bold) and between (outside diagonal) clusters based on the dissimilarity matrix obtained by Kohonen’s Self-Organizing Maps.

| Cluster | I | II | III | IV | V |
| --- | --- | --- | --- | --- | --- |
| I | **0,261** | 0,684 | 0,990 | 0,986 | 1,000 |
| II | - | **0,271** | 0,991 | 0,873 | 0,993 |
| III | - | - | **0,352** | 0,937 | 0,734 |
| IV | - | - | - | **0,232** | 0,707 |
| V | - | - | - | - | **0,258** |

When comparing clusters, it is verified that the greatest divergence was established between cluster I and V, with genetic distance between them equal to 1.00. The graphical dispersion of the dissimilarity matrix shown in Fig 5 shows that the dispersion between and within the clusters was high, and that cluster I was very distant from clusters III, IV and V. It is also possible to verify that families F8 and F23 were positioned very close to cluster I (Fig 5), but not enough to be grouped in cluster I by the UPGMA method (Fig 3). This was probably due to the presence of some characteristic strongly associated to the other genotypes that compose the clusters II. In relation to the efficiency of the clusters using the Anderson discriminant analysis (Table 2), it was verified that the classification adopted allowed a rate of 100% accuracy.

**Table 2.** Percentage of correct classifications by the Anderson discriminant function for each cluster formed by the UPGMA method.

| Cluster | I | II | III | IV | V |
| --- | --- | --- | --- | --- | --- |
| I | 100 | 0 | 0 | 0 | 0 |
| II | 0 | 100 | 0 | 0 | 0 |
| III | 0 | 0 | 100 | 0 | 0 |
| IV | 0 | 0 | 0 | 100 | 0 |
| V | 0 | 0 | 0 | 0 | 100 |

**Figure 5.**
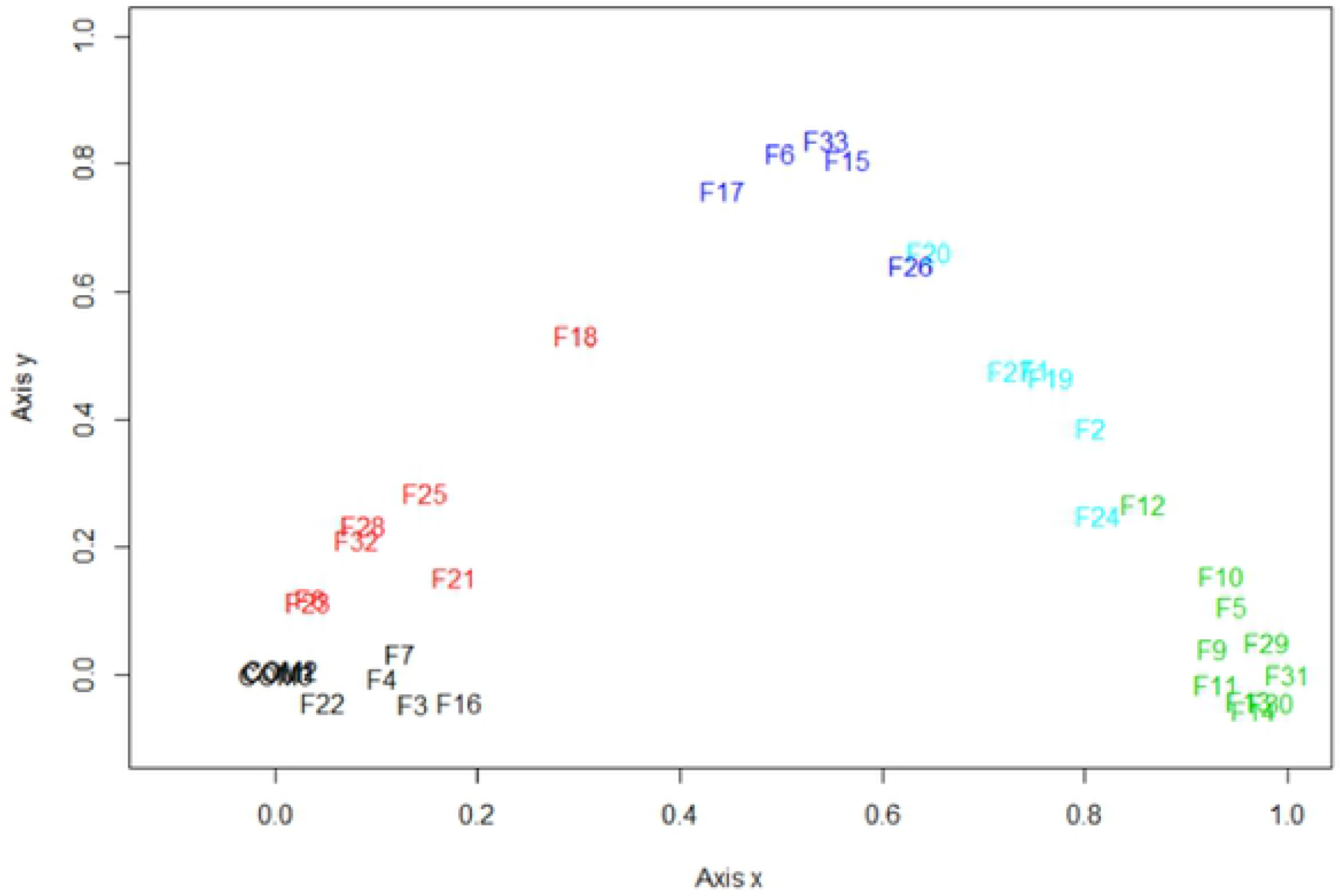
Graphic dispersion of the dissimilarity matrix in 1000 SOM type networks with five rows and one column (5 neurons). The colors represent the four distinct clusters formed: 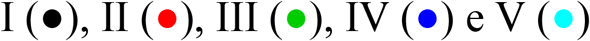.

## Discussion

The efficiency of the use of neural networks in breeding programs has been presented in several studies, such as the selection of more productive genotypes of sugarcane (*Saccharum* spp.) [19] and the evaluation of genetic divergence (*Vitis vinifera* L.) [20], papaya (*Carica papaya* L.) [1] and guava (*Psidium guajava* L.) [9].

In kale the neural networks were used by Azevedo et al. [11] in leaf area prediction. The technique proved to be feasible to estimate the leaf area and to assist in the selection of superior genotypes. However, there are no other studies carried out with the crop and RNAs, which reinforces the importance of the study for application in breeding, especially for the study of genetic divergence.

The self-organizing maps began to be explored by T. Kohonen as of 1982, however their use is becoming more frequent [13]. The SOM-type networks present unmasked clustering process and perform the training based on the patterns of their spatial organization only in the homology (similarity) of the data, that is, without prior knowledge of the class to which they belong [21].

This allows the technique to be an appropriate procedure for grouping the genotypes based on morphoagronomic characters, since there is no requirement for calibration [22] without the need for information on experimental designs and genealogies, which in some situations is unknown. The high divergence observed in the matrix of dissimilarity (Fig 2) and the dispersion of the families in the graphical representation (Fig 5) is indicative of genetic variability in the population, which points out its potential for genetic improvement [8]. Significant genetic gains with selection are conditioned by the existence of such genetic variability [23].

The representation of the matrix by the dendrogram (Fig 3) also showed a high co-optic correlation, which shows high agreement in the grouping performed and the dissimilarity matrix [24]. This reinforces the reliability of subsequent crossing to be established based on the composition of the clusters using the RNAs. González-Cuéllar and Obregón-Neira [25] also point out that the main advantage associated with Kohonen’s self-organizing maps is the easy visualization of the established classification, since in other classification methods involving a greater number of variables the process becomes more complex.

Although it does not present families of high yield, cluster I is composed of materials with characteristics of interest for the improvement of the kale, since it resembles the commercial cultivars. Among these characteristics, the smaller petiole length, the lower plant height and the stem, and especially the low number of shoots, stand out. This approach is closer to what is required as a commercial and agronomic standard, because in addition to resistance to the main pests, the genetic improvement of kale must prioritize genotypes with lower height, fewer shoots and more commercial leaves. This favors higher yield, facilitates crop management, harvesting, and reduces costs with plant staking. Thus, it is indicated to include these families to compose the recombinant population, along with other families of higher yield and number of leaves, as observed for the families of clusters III and V.

Another relevant observation is that for most of the families evaluated in this study, the production and number of leaves presented a positive degree of association with the increase in the number of shoots, that is, more productive families tended to have a higher number of shoots. This differs from Azevedo et al. [26], who found a low degree of association between the number of shoots and the mass of fresh leaves when evaluating genetic correlations in leaf cabbage genotypes.

This association identified in the study population is not of interest for the process of improvement of the same. This can favor the vegetative propagation, which generates higher cost with crop by the producer. The presence of this characteristic also reduces the producer’s dependence on seeds, which may not be favorable when commercial cultivars are to be obtained. Therefore, it is necessary that future recombinations prioritize the crossing of low sprouting plants with high yield plants.

The reduction of the number of shoots can be done from crossing between the clusters I families (low plants and with few buds) with those belonging to clusters III and V, which presented high yield, greater number of leaves and larger stem diameters. These clusters were also genetically distant (Table 1 and Fig 5). This is important in the recombination process, since it increases the probability of obtaining superior genotypes in yield and with agronomic characteristics of interest in the segregating population. Important crosses can also be established between the families of clusters II with families of clusters III and V, since they are the most genetically distant.

The genetic distance between clusters and within clusters (Table 2) and graphical dispersion (Fig 5) show that although the UPGMA method forms clusters with families of similar characteristics, they present high variability among themselves, and also within clusters formed. This corroborates with that observed in the dissimilarity matrix formed from Kohonen’s self-organizing maps (Fig 2), which indicated high divergence among genotypes. This can occur because the population is in the early stages of breeding.

As for the efficiency of the adopted methodology, Anderson’s discriminant analysis showed that it is adequate and efficient. This is in agreement with Barbosa et al. [1], which highlights that the discriminant analysis is useful in checking the consistency of clusters, and is recommended for this type of study. Thus, it is clear the consistency of the methodology used and its indication for future work.

## Conclusions

Crossings between families belonging to clusters I and families of clusters III and V are the most recommended to allow the selection of plants with lower number of shoots, plant height, leaf length, leaf width, petiole length and greater number of leaves, leaf yield and stem diameters.

The proposed methodology for the use of SOM networks is efficient, since it allows the selection of the network configuration without subjectivity and high rate of correctness by the Anderson descending analysis, forming clusters in agreement with the morphological data.

## Supporting information

**S1 Table.** Database for variables evaluated in half-sib families progenies of leaf kale. (xlsx)

**S2 Table.** Script programming. (DOCX)

## Acknowledgment

To CNPq, FAPEMIG and CAPES for the granting of scholarships and financial resources for the development of the project. This study was financed in part by the Coordenação de Aperfeiçoamento de Pessoal de Nível Superior - Brasil (CAPES) - Finance Code 001. To UFVJM for the opportunity to develop the study and germplasm supply.

## Author Contributions

**Conceptualization:** Orlando Gonçalves Brito, Valter Carvalho de Andrade Júnior, Alcinei Mistico Azevedo.

**Data Curation:** Orlando Gonçalves Brito, Alcinei Mistico Azevedo.

**Formal Analysis:** Orlando Gonçalves Brito, Alcinei Mistico Azevedo.

**Investigation:** Orlando Gonçalves Brito, Maria Thereza Netta Lopes Silva, Ludimila Geiciane de Sá, Clóvis Henrique Oliveira Rodrigues, Ana Clara Gonçalves Fernandes.

**Resources:** Valter Carvalho de Andrade Júnior.

**Writing – Original Draft Preparation:** Orlando Gonçalves Brito, Valter Carvalho de Andrade Júnior, Alcinei Mistico Azevedo.

**Writing – Review & Editing:** Orlando Gonçalves Brito, Alcinei Mistico Azevedo, Valter Carvalho de Andrade Júnior.

